# Hi-GREx: A 3D Genome–Guided Framework for enhancing Gene Expression Prediction Using Hi-C–Selected Distal SNPs

**DOI:** 10.1101/2025.09.26.678794

**Authors:** Ketki Joshi, Min Chen, Zhenyu Xuan

**Author notes:** Corresponding author; Correspondence to Zhenyu Xuan.

## Abstract

Genome-Wide Association Study (GWAS) method has been successfully used to map thousands of loci associated with complex traits, but its ability to reveal the molecular mechanisms altered in complex diseases has been limited due to not including combinations and interactions between markers when predicting a disease. Transcriptome-Wide Association Studies (TWAS) estimate the aggregate effects of multiple genetic variants on complex diseases and represent a promising approach to address the limitations of GWAS. In particular, TWAS provides insights into the functional consequences of disease-associated SNPs by linking them to gene transcription, thereby offering a mechanistic understanding that GWAS alone cannot provide.

However, TWAS associated variants have been annotated with the closest or most biologically relevant candidate gene within arbitrarily defined distances but fails to account for long distance SNPs which can affect many genes and have a widespread impact on regulatory networks. Therefore, there is a need to leverage these observed enrichments and build a method that incorporates both short and long distance-associations between SNPs and complex phenotypes.

Here we present a method which can utilize Hi-C data to capture “informative” long-distance SNPs and aim to improve prediction accuracy of previous TWAS method.

We benchmarked our method on GTEx brain cortex genotype and expression data together with the corresponding Hi-C data. By using the “informative” long distance SNPs selected based on Hi-C, our method improved prediction accuracy of gene expression for 77.4% of the active genes across the entire genome. Particularly, our method can build significant expression models for 18% of genes which were missed by using only short-distance SNPs. Our method has demonstrated the efficiency and importance of utilizing long-distance SNPs in predicting gene expression and can further enhance the power of TWAS methods.

## Introduction

Expression quantitative trait loci (eQTLs) provide a critical framework for elucidating the molecular mechanisms by which genetic variants regulate gene expression and contribute to complex traits and diseases [1–4]. By identifying genetic loci that modulate transcript abundance, eQTL mapping has advanced the interpretation of non-coding variants—many of which are enriched among genome-wide association study (GWAS) signals—and has facilitated the prioritization of candidate genes at trait-associated loci [4,5]. Despite these advances, eQTL studies remain predominantly focused on cis-regulatory variants—those located within a 1Mb window of the gene—due to their relatively strong effect sizes and statistical tractability [6].

To better connect regulatory variation to complex traits, transcriptome-wide association studies (TWAS) have been developed. Among these, the PrediXcan framework has gained widespread adoption for integrating genetically predicted gene expression with GWAS summary statistics [7]. PrediXcan constructs predictive models using elastic net regression on cis-eQTLs to impute gene expression levels in relevant tissues, which are then tested for association with complex phenotypes. This gene-centric approach increases statistical power, reduces the burden of multiple testing, and improves the interpretability of association signals relative to single-variant GWAS. The FUSION method extends this strategy by leveraging multiple modeling approaches, including best linear unbiased prediction (BLUP), Bayesian sparse linear mixed models (BSLMM), LASSO, and elastic net, allowing it to flexibly capture diverse genetic architectures underlying gene expression [37]. UTMOST further refines prediction by performing penalized multivariate regression across multiple tissues, enabling the detection of shared cross-tissue regulatory effects and improving prediction accuracy for genes expressed in multiple contexts [38]. TIGAR employs a Bayesian Dirichlet process regression framework, which reduces reliance on parametric assumptions and is particularly useful for genes with low expression heritability [39]. In addition, polygenic risk score (PRS)-based approaches aggregate the effects of multiple variants genome-wide to predict gene expression or complex traits [40,41]; while these methods can capture broader genetic variation, they typically do not incorporate explicit regulatory mechanisms or distal interactions, limiting biological interpretability.

Nevertheless, the power of PrediXcan and related gene-based prediction approaches is currently limited by their focus on cis-regulatory variation—those variants physically close to the gene [6]. This restriction exists largely for computational and statistical reasons, as cis-eQTLs are easier to detect, but mounting evidence suggests that trans-acting variants (which regulate genes on different chromosomes or at significant genomic distances) may explain a substantial portion of the heritability of gene expression and complex traits [8]. For example, studies in twins have shown that cis-SNPs may explain as little as 10% of gene expression variation in some tissues [8]. Critically, some GWAS risk variants function via long-range looping mechanisms, exemplified by the obesity-associated FTO locus, which regulates IRX3 via distal chromatin interaction, rather than the nearest gene [9,10].

The challenge is heightened by the fact that the vast majority of disease-linked variants identified by GWAS reside in non-coding regions, where their target genes and mechanisms are often unclear [10]. Analyzing all possible distant regulatory relationships genome-wide is both computationally intensive and statistically challenging, motivating the adoption of strategies that can efficiently prioritize “informative” long-range variant-gene interactions. This is where emerging three-dimensional chromatin conformation capture technologies become vital. Hi-C provides a genome-wide map of physical DNA–DNA contacts, thereby revealing direct evidence of regulatory interactions that may not be predictable from linear proximity [11–13]. By informing which distal regions are likely to affect gene regulation, Hi-C data can guide the selection of candidate variants for feature selection and help close the gap between GWAS signals and the true molecular drivers of disease. By mapping chromosomal interactions at a genome-wide scale, Hi-C offers a unique opportunity to interpret non-local gene regulation, enriching our understanding of genetic architecture in health and disease.

We propose a novel method, **Hi-GREx**, which integrates both proximal (short-distance) and distal (long-distance) SNPs—encompassing both coding and non-coding regions— to construct gene expression prediction models. Proximal SNPs, located within a 1Mb window of the gene, directly regulate gene expression and are typically included in conventional models. In contrast, distal SNPs can modulate the expression of target genes either through direct or indirect mechanisms. Direct regulation can occur via chromatin looping, which brings distant regulatory elements into close spatial proximity with gene promoters in three-dimensional (3D) nuclear space. Alternatively, distal SNPs may exert indirect (mediation) effects, for instance via a mediator gene; here, a cis-regulatory SNP influences its proximal gene, which in turn transmits the regulatory effect to a distal gene—manifesting as trans-eQTL associations [14–17] [Figure 1A].

**Figure 1:**
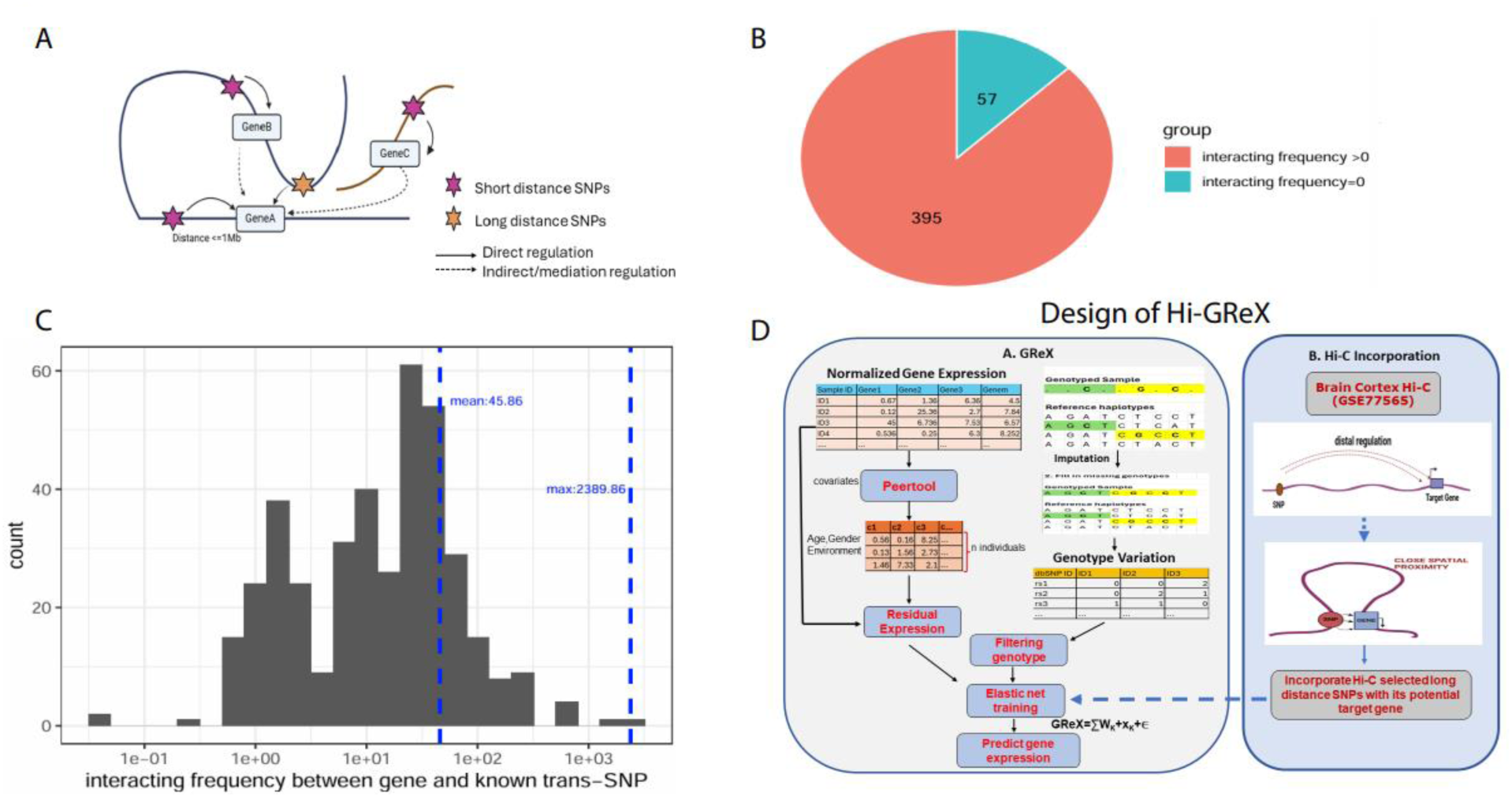
(a) Schematic of Gene Regulation by SNPs. Gene regulation may occur either via short-distance SNPs—located within 1Mb of the target gene—or via long-distance SNPs, which exert their effects through chromatin looping in three-dimensional nuclear space. Direct gene regulation is indicated with solid arrows, while dotted arrows denote indirect or mediated genetic effects on gene expression. (b) Pie Chart: Depicts the proportion of intrachromosomal trans-eQTLs supported by chromatin interaction evidence. *(c)* Histogram: Shows the distribution of chromatin interaction frequencies for intrachromosomal trans-eQTLs, capturing the diversity in interaction strength among these regulatory events. *(d)* Workflow for Incorporating Chromatin Interaction Data into Gene Expression Models. The left panel (A) outlines the standard GReX modeling framework: Reference transcriptome data (including genotypes and gene expression from brain cortex tissue, as provided by GTEx) form the basis for training gene expression prediction models. Genotype data are imputed and quality controlled using criteria for minor allele frequency (MAF) and imputation quality (R²). Covariates capturing non-genetic influences—including demographic, environmental, and hidden factors—are regressed out to obtain normalized residual expression levels. A regularized additive model, implemented via elastic net regression, is then fit to predict gene expression from SNP genotype data, constituting the conventional TWAS approach, which typically restricts input SNPs to those nearby each gene. The right panel (B) expands this framework by integrating Hi-C–derived chromatin interaction data. Hi-C datasets for brain cortex tissue, obtained from NCBI, are used to identify “informative” long-distance SNPs physically interacting with target genes via chromatin looping. These distal SNPs, brought into spatial proximity with their regulatory targets, are incorporated as model features to re-train gene expression predictors and assess whether their inclusion augments TWAS detection power.

Our central hypothesis is that incorporating Hi-C-guided distal SNPs—in addition to conventional proximal SNPs—significantly enhances the accuracy and robustness of gene expression prediction models. Unlike existing approaches that rely solely on nearby cis-variants, Hi-GREx leverages three-dimensional chromatin interaction data to identify distal variants that physically contact gene promoters and are thus more likely to play a regulatory role. By selectively incorporating these biologically informed, spatially distal SNPs as informative features, Hi-GREx enriches the predictive landscape of gene expression models, capturing regulatory effects that would otherwise be missed in linear analyses. This expanded feature set improves model performance by explaining a greater proportion of gene expression variability, ultimately increasing the power to detect trait-associated loci. Our results demonstrate that Hi-GREx consistently outperforms conventional gene-based prediction methods across diverse genetic architectures, highlighting the utility of integrating chromatin conformation data to build more comprehensive and biologically meaningful expression models.

## Results

### Chromatin Interactions Reveal Physical Links Between Known trans-eQTL and Target Genes

To investigate the physical relationships between known trans-eQTL and their target genes, we analyzed published trans-eQTL data from Klein N. et al [27]. Focusing on *intrachromosomal* trans-eQTL pairs, we leveraged Hi-C chromatin interaction data, restricting our analysis to interactions occurring on the same chromosome due to the limitations of available intrachromosomal Hi-C datasets. Importantly, all trans-eQTL analyzed were separated by more than 1 Mb from their target gene, which aligns with our aim to explore distal regulatory variants positioned beyond conventional cis-eQTL search windows.

Our findings demonstrate that a substantial proportion (87.38%) of the known intrachromosomal trans-SNPs show direct chromatin interactions with their respective target genes [Figure 1B]. This high overlap supports the hypothesis that long-range chromatin interactions frequently mediate the regulatory effects of distal trans-SNPs. The remaining trans-eQTL without detected chromatin contacts may represent genuine biological exceptions or could be artifacts stemming from incomplete coverage or loss of regions during the genomic coordinates liftover process. Further, analysis of the chromatin interaction frequencies between known trans-SNPs and their target genes revealed a wide distribution, with interaction counts ranging up to a maximum of 2,389 [Figure 1C]. The mean interaction frequency was observed at 45.86, underscoring the variability in the physical contact between trans-eQTL and gene targets, as expected for regulatory elements that act over large genomic distances.

### Optimized SNP Selection and Chromatin-Guided Integration Improve Gene Expression Prediction

Effective feature selection is critical in high-dimensional genotype datasets to reduce dimensionality, minimize redundancy, and prevent model overfitting. In this study, we utilized the matrix-eQTL approach (p < 0.05) [18] to identify SNPs significantly associated with gene expression and systematically varied the number of input features per gene (ranging from 50 to 250), constrained by a cohort of 205 brain cortex samples.

Model performance evaluated through Spearman correlation (Rho) between predicted and residual gene expression, reached its peak at 150 input SNPs per gene (mean Rho = 0.51). Increasing the number of features beyond this threshold resulted in diminishing predictive returns, with average correlations decreasing to Rho = 0.48 and Rho = 0.47 for 200 and 250 SNPs [Figure S1], respectively, likely due to the inclusion of redundant or weakly informative features. These results underscore the importance of rigorous feature selection and suggest that, for datasets of comparable size, incorporating approximately 150 top-associated SNPs per gene provides an optimal trade-off between predictive power and model complexity. This finding underscores that larger numbers of input features do not necessarily translate into improved prediction and may, in fact, impair model performance.

To maximize the predictive accuracy of gene expression using genotype data, we systematically evaluated several strategies for incorporating long-distance SNPs alongside proximal SNPs. We hypothesized that distal SNPs identified both as eQTLs and as physical interactors with the gene promoter in Hi-C data—thus “informative” by both association and 3D nuclear context—would provide the most value for expression prediction.

To test this, we compared the performance of six modeling strategies:

1. **AllShort:** Using all available short-distance (<1 Mb) SNPs with no feature selection.
2. **150Short:** Using only the top 150 short-distance SNPs ranked by association strength.
3. **RandomLong:** Using n modeled short-distance SNPs (n varies per gene), with the remaining (150-n) features filled with randomly selected long-distance (>1 Mb) SNPs.
4. **HiCselected:** Using n modeled short-distance SNPs, with (150-n) filled by long-distance SNPs supported by chromatin (Hi-C) interactions with the gene, regardless of eQTL significance.
5. **eQTLselected:** Using n modeled short-distance SNPs, and (150-n) filled by long-distance SNPs chosen solely on eQTL association (p-value) without chromatin interaction support.
6. **HiCselected-eQTLranked (HiGReX):** Using n modeled short-distance SNPs and (150-n) long-distance SNPs that are both significant eQTLs and supported by Hi-C chromatin interactions.

Models using feature selection among only the short-distance SNPs (150Short) outperformed models using all available short-distance SNPs (AllShort), with mean Spearman correlation improving from 0.51 to 0.53. This supports feature selection as a key step to reduce redundancy and noise. Integration of long-distance SNPs randomly (RandomLong) or solely by Hi-C or eQTL criteria did not lead to improved predictive performance, showing mean correlation values of 0.49 (RandomLong), 0.49 (HiCselected), and 0.50 (eQTLselected). These approaches likely introduce non-informative or redundant SNPs that dilute predictive power [Figure 2A].

**Figure 2:**
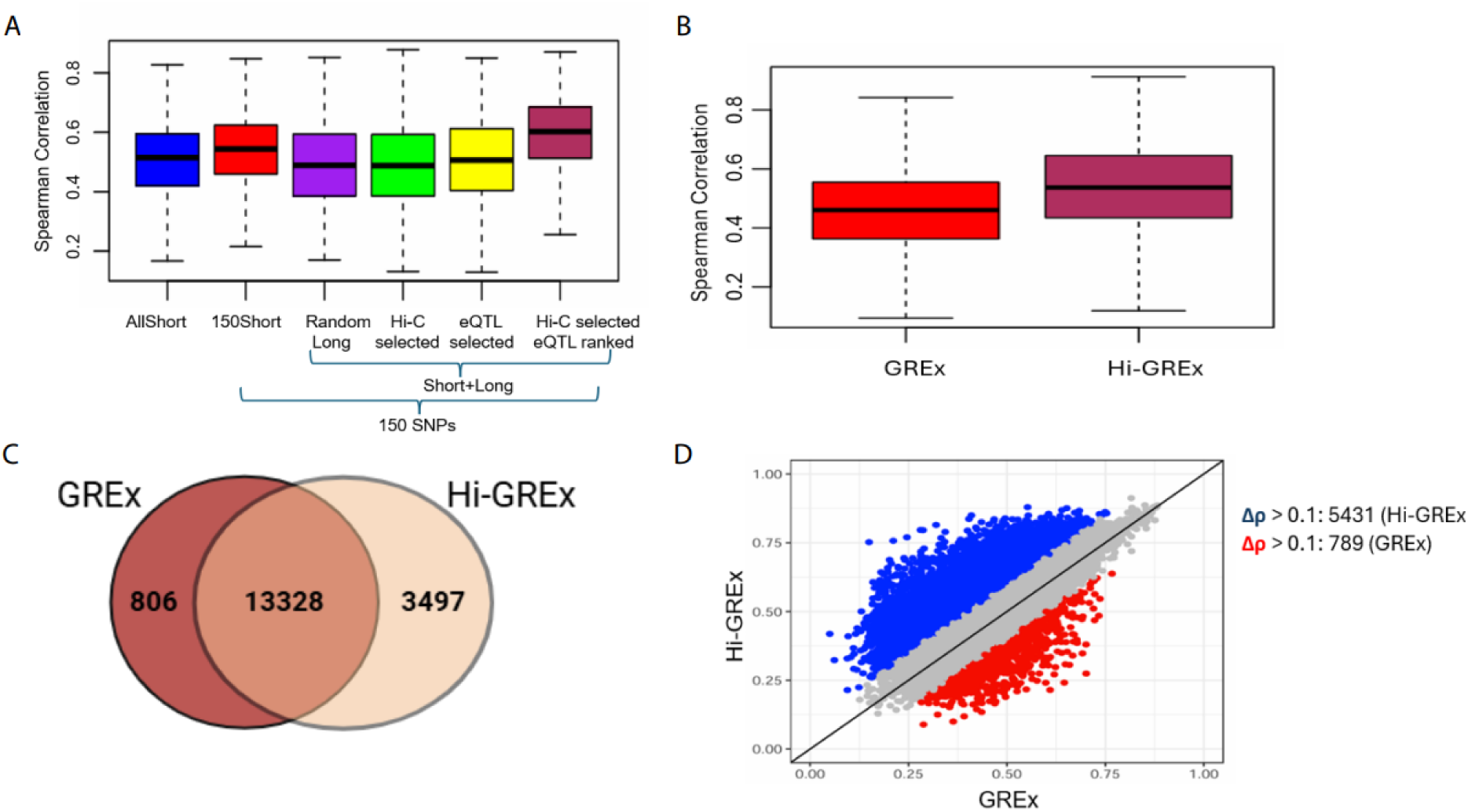
**(a)** Boxplot showing the distribution of spearman correlation between predicted and residual expression for genes in chr(1,2,3,20,21,22) (number of genes=1398) across different models. Long distance SNPs are incorporated with short distance SNPs based on different criteria (i) random selection (ii) Hi-C selection (iii) eQTL selection (iv) Both Hi-C selection and eQTL ranking. **(b)** Boxplot showing the distribution of correlation between predicted and residual expression for GREx vs Hi-GREx for all the genes across the genome (n=13328). An improvement is observed after including Hi-C selected “informative” long distance SNPs in comparison to GREx models **(c):** Venogram showing count of gene models which are unique to each case and common genes being modelled by all the methods. (d) Scatter plot showing spearman correlation for each gene across different models [GREx, Hi-GREx], a greater number of genes have correlation higher than 0.1 after including long-distance SNPs.

Remarkably, the **HiCselected-eQTLranked (HiGReX)** approach—systematically selecting long-distance features that are both physically proximal in 3D genome space (by Hi-C) and significant as eQTLs—led to the best performance with a mean Spearman correlation of 0.59. For 669 out of 1,398 modeled genes (47.85%), this strategy outperformed all other methods, including those using only local SNPs [Figure 2A].

### Integration of Chromatin-Informed Distal Variants Enhances Transcriptome Prediction Across the Genome

We systematically quantified the impact of incorporating Hi-C–selected distal SNPs into transcriptome prediction models. Specifically, we constructed GREx models and compared them to HiGREx model for the GTEx brain cortex dataset (n = 205). Model performance was evaluated based on both the number of genes for which expression could be robustly predicted and the accuracy of these predictions, as measured by the Spearman correlation between observed and predicted expression levels.

Among 18,916 genes analyzed, 13,328 were successfully modeled by both approaches. Across this shared set, integration of spatially prioritized distal SNPs via the Hi-GREx approach yielded substantial improvements in predictive power. Notably, 74.9% (9,995/13,328) of modeled genes exhibited higher predictive accuracy with Hi-GREx relative to conventional GREx (mean Spearman’s ρ: 0.51 vs. 0.45) [Figure 2B]. Moreover, 5431 genes (40.7%) experienced a marked increase in model fit (Δρ > 0.1), whereas only 789 genes demonstrated such an increase for GREx over Hi-GREx. Visualization via scatterplot reveals a predominant shift above the diagonal, indicating widespread and often pronounced gains in accuracy with Hi-GREx [Figure 2D]

The Venn diagram further highlights the broadening of transcriptome coverage afforded by chromatin-guided distal variant prioritization: 3,497 genes could be modeled exclusively by Hi-GREx—constituting 19% of all genes modeled by this approach— while only 806 were uniquely captured by GREx [Figure 2C].

To further evaluate the performance of Hi-GREx across tissues, we compared its predictive power with the conventional GREx model in spleen, pancreas, and brain cerebellum [Table 1]. In all three tissues, Hi-GREx consistently improved prediction accuracy relative to GREx, as reflected by higher mean R² values (0.49 vs. 0.45 in spleen, 0.52 vs. 0.46 in pancreas, and 0.58 vs. 0.49 in brain cerebellum). Importantly, Hi-GREx also substantially increased the number of genes with predictive models, adding ∼2,000–2,500 genes beyond those captured by GREx in each tissue. These improvements demonstrate that Hi-GREx method of incorporating Hi-C selected long-range SNPs enhances the predictive capacity of gene expression models in a manner that is both robust and generalizable across different tissues

**Table 1.** Model performance comparison of GREx and Hi-GREx across tissues. The table shows the performance of genetically regulated expression (GREx) and hierarchical GREx (Hi-GREx) models across three tissues (spleen, pancreas, and brain cerebellum). For each tissue, the sample size, number of genes with predictive models, and mean rho values are reported. Hi-GREx consistently increases the number of modeled genes and achieves higher mean rho compared to GREx, indicating improved prediction accuracy.

| **Model Performance for GREx vs Hi-GREx Across Tissues** |  |  |  |  |
| --- | --- | --- | --- | --- |
|  |  | **Sample Size** | **Genes** | **Mean R<sup>2</sup>** |
| **Spleen** | GREx | 227 | 11365 | 0.45 |
|  | Hi-GREx | 227 | 14532 | 0.49 |
| **Pancreas** | GREx | 305 | 11650 | 0.46 |
|  | Hi-GREx | 305 | 14302 | 0.52 |
| **Brain Cerebellum** | GREx | 209 | 14818 | 0.49 |
|  | Hi-GREx | 209 | 17222 | 0.58 |

### Higher Chromatin Interaction Observed for Modelled Long-Distance SNPs Compared to Input Feature Sets

We assessed chromatin interaction frequencies between genes and long-distance SNPs selected in the final HI-GREx model, comparing these to long-distance SNPs not selected by the model but included as input features. Our underlying hypothesis is that long-distance SNPs prioritized by the model should demonstrate stronger chromatin interaction with their associated genes than background SNPs not prioritized in the model.

We evaluated the cumulative distribution functions (CDFs) for three groups: known trans-SNPs (blue curve), model-selected long-distance SNPs (green curve), and input long-distance SNPs not selected by the model (red curve). The CDF for known trans-SNPs is notably shifted toward higher interaction frequencies, consistent with the expectation that these represent true regulatory connections. Importantly, the model-selected long-distance SNPs also show a rightward shift (toward higher interaction frequencies) compared to non-selected input SNPs, supporting the model’s ability to capture SNPs with stronger chromatin connectivity. In contrast, non-selected input SNPs display the lowest interaction frequencies, indicating weaker or absent chromatin interactions at a distance [Figure 3A].

**Figure 3:**
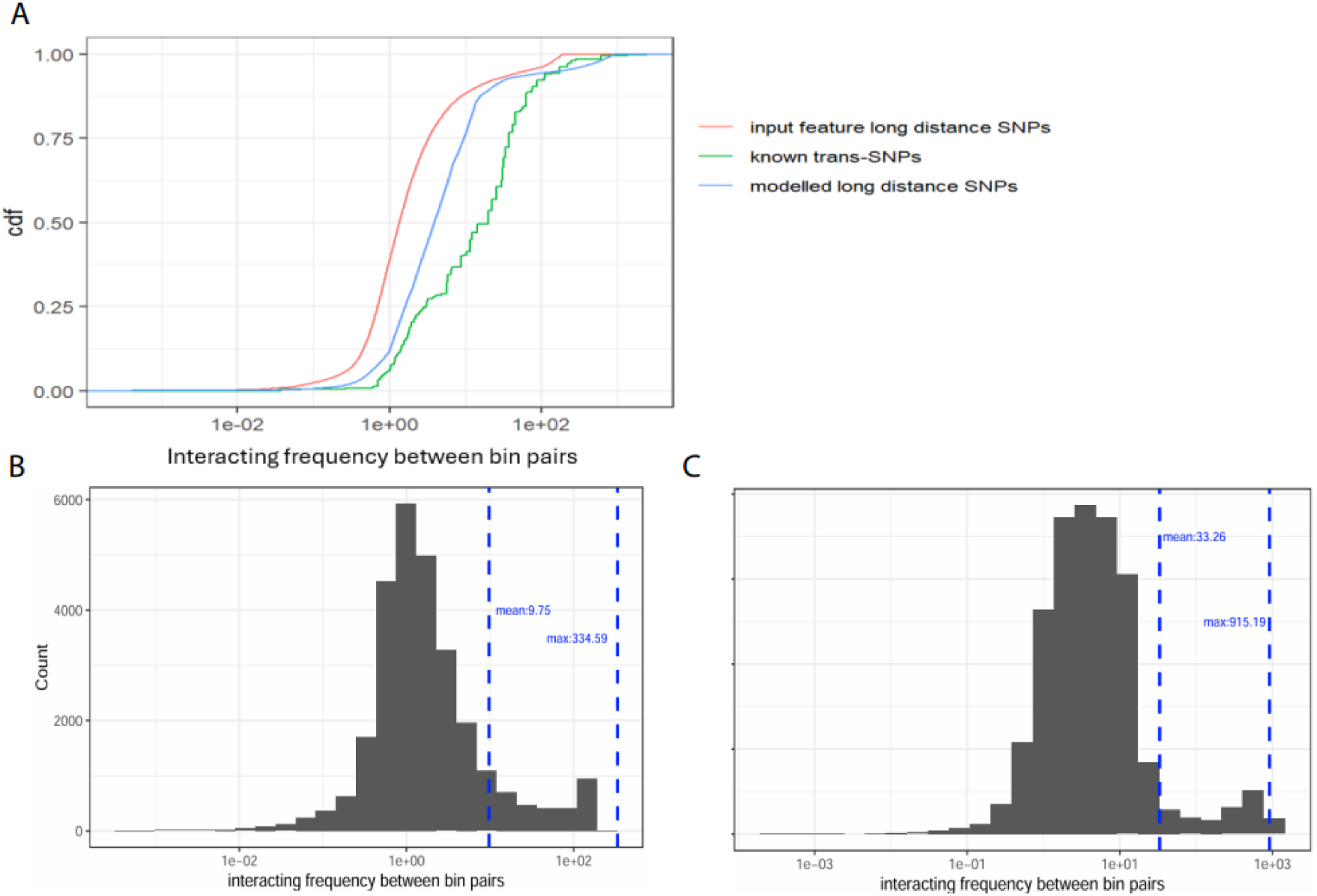
**(a)**: Cumulative distribution function plot for comparing the frequency of interaction between gene and SNP bin pair for known trans-SNPs vs modelled long distance SNPs and input feature-long distance SNPs. **(b):** Histogram showing distribution of chromatin interaction frequency for long distance SNPs selected present in input feature set but not selected in final Hi-GREx model **(c):** Histogram showing distribution of chromatin interaction frequency for long distance SNPs selected in final Hi-GREx model having much higher interacting frequency in comparison to not modelled features

Quantitative comparisons reinforce these findings. As shown in Figure 3C, the mean and maximum chromatin interaction frequencies between genes and modelled long-distance SNPs closely match those observed for known trans-SNPs [Figure 1C], with mean values around 33.26 and maximum values reaching over 900. Some model-selected SNPs approach interaction frequencies near 1000, underscoring their strong regulatory potential. In contrast, non-modelled background long-distance SNPs exhibit substantially lower mean (9.75) and maximum (334.59) values with no bins reaching interaction frequencies of 1000 or higher, as depicted in Figure 3B.

Collectively, these data support the conclusion that the HI-GREx model preferentially selects long-distance SNPs exhibiting robust chromatin interactions with their target genes, congruent with established trans-eQTLs. This enrichment is not observed in the broader set of input SNPs not prioritized by the final model, indicating the added biological relevance of model selection criteria.

### Integrating Long-Range Genetic Interactions Dramatically Improves Predictive Modeling of Alzheimer’s-Associated Gene Expression

We observe substantial improvement in the accuracy of predicted gene expression for key Alzheimer’s disease (AD)-associated genes—PSEN1, PSEN2, APP, and ADAM10 for Hi-GREx case. Figure 4 A compares the performance of two modeling approaches. On the left, the GREx model incorporates only proximal SNPs, often resulting in distinct vertical striations that indicate predicted expression is disproportionately driven by a small number of variants. On the right, the Hi-GREx model integrates additional long-range (chromatin interaction–derived) SNPs, providing a more continuous and distributed contribution from multiple loci. This is evident from the pronounced increase in the correlation coefficients (rho values) across all four genes shown, most notably for PSEN2 (from 0.34 to 0.53) and PSEN1 (from 0.20 to 0.41). We also provide a list of 50 AD associated genes and their correlation developed by using GREx vs Hi-GREx case and their delta correlation [Comparison_of_Performance_ADGenes.xlsx]. Figure 4B iillustrates the above concept using APP as an example. The schematic diagrams show that including long-range SNPs (such as rs79345946 and rs80269739, with strong effect sizes) in the Hi-GREx model not only deepens the mechanistic interpretation of regulatory effects but also redistributes the overall contribution to gene expression across multiple regulatory elements, as seen in the scatterplots below the diagrams. Consequently, the predictive correlation for APP expression jumps from 0.17 with traditional modeling to 0.41 in the integrated model. These findings highlight how leveraging three-dimensional genome architecture greatly refines expression prediction for genes implicated in AD, where genetic regulation is often polygenic and dispersed across the genome.

**Figure 4:**
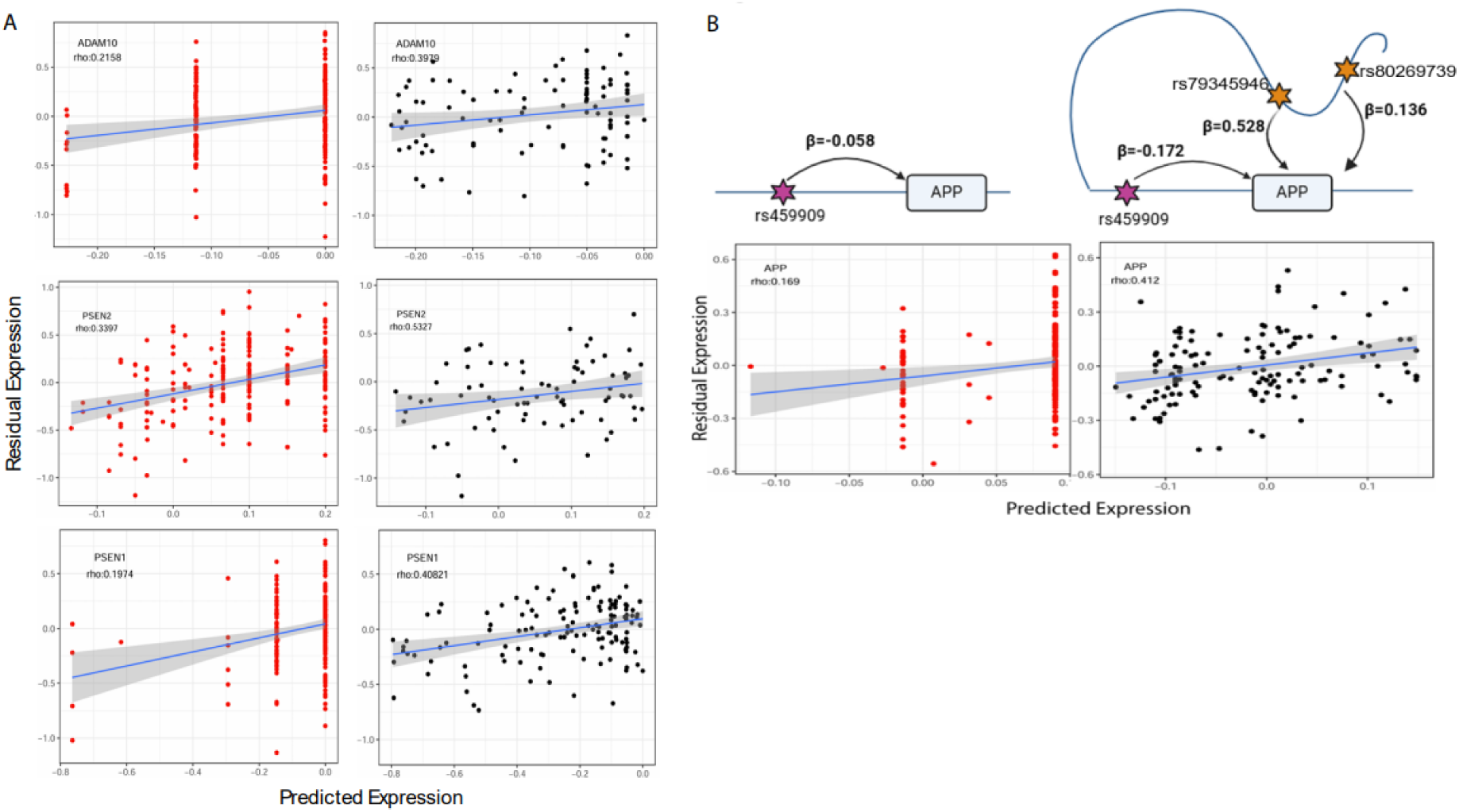
**(a)** These plots show residual vs. predicted levels for 3 well predicted AD genes. Weights stored in GTEx brain cortex elastic net models were used to compute predicted levels. (**b**) beta coefficient of the common GREx SNP rs459909 in APP gene model (i) For GREx model only. (ii) For Hi-GREx model, beta coefficient of rs459909 is increased in the presence of long-distance SNPs. Scatter plot for APP, represented SNPs in each model with their rho values, a higher prediction is observed in the presence of Hi-C selected long distance SNPs “rs79345946” and “rs80269739”

### Application of Hi-GREx to AD

We examined whether transcriptional prediction models incorporating three-dimensional chromatin interaction data (via Hi-C) can prioritize functionally important genetic associations in Alzheimer’s disease (AD), compared to traditional expression prediction approaches limited to local regulatory variants. To address this, we applied S-PrediXcan [Supplementary_document.txt] to brain cortex tissue using multiple expression prediction models, including both conventional short-range SNP models (GREx) and those incorporating Hi-C–informed distal SNPs (Hi-GREx). Summary statistics were derived from a recent large-scale AD genome-wide association study (GWAS) [28]. Application of the GREx models yielded 30 significant gene associations with AD at a Bonferroni-corrected threshold (p < 0.05; **Figure 5A**). In contrast, integrating long-range chromatin contacts via Hi-GREx substantially increased the statistical power, identifying 188 significant AD-associated genes (**Figure 5B**). Furthermore, comparison with the established ADSP GVC gene list [36] revealed that Hi-GREx models resulted in a greater overlap with known AD risk genes than conventional GREx models (**Figure 5C** demonstrating that the inclusion of distant regulatory SNPs increases sensitivity for pathogenic gene discovery. Supporting this, the genes SORL1, CR1, and TREM2—all well-established in AD pathogenesis—showed markedly more significant associations in the Hi-GREx model compared to conventional GREx. For example, SORL1 (Hi-GREx *p* = 7.35 × 10⁻ ²⁰ vs. GREx *p* = 2.34 × 10⁻ ⁶) is a key regulator of amyloid precursor protein (APP) trafficking and has been strongly implicated in both early- and late-onset AD [31]. CR1 (*p* = 4.48 × 10⁻ ²² vs. 2.04 × 10⁻ ¹⁶) is involved in the complement cascade and immune clearance of amyloid-β, and its genetic variants have been associated with AD susceptibility in multiple GWAS studies [32,33]. Similarly, TREM2 (*p* = 8.51 × 10⁻ ⁸ vs. 3.27 × 10⁻ ⁶) plays a critical role in microglial activation and response to amyloid pathology, and rare coding variants in this gene confer a significant increase in AD risk [34,35]. The increased statistical significance for these genes in the Hi-GREx model suggests that their expression may be regulated by distal enhancers not captured by traditional local approaches [S-PrediXcan_AssociationResults.xlsx]. We also found examples where SNPs identified through GWAS were originally assigned to their nearest genes based on genomic proximity. However, analysis using the HiGReX model revealed that these SNPs are located within the regulatory architecture of distal genes. These model genes, rather than the nearest genes reported by GWAS, are the likely functional targets influenced by the SNPs [Table S2]. Importantly, the SNPs are included in the gene models not due to proximity but because they fall within regulatory elements—such as enhancers—that interact with the promoters of these distal genes. This indicates that the association of these SNPs with Alzheimer’s disease (AD) risk may be mediated through regulation of these distal model genes, rather than through the nearest genes initially reported. These findings underscore the limitations of proximity-based SNP-to-gene assignments in GWAS and highlight the utility of integrative models like HiGReX for uncovering more accurate regulatory relationships underlying complex traits.

**Figure 5:**
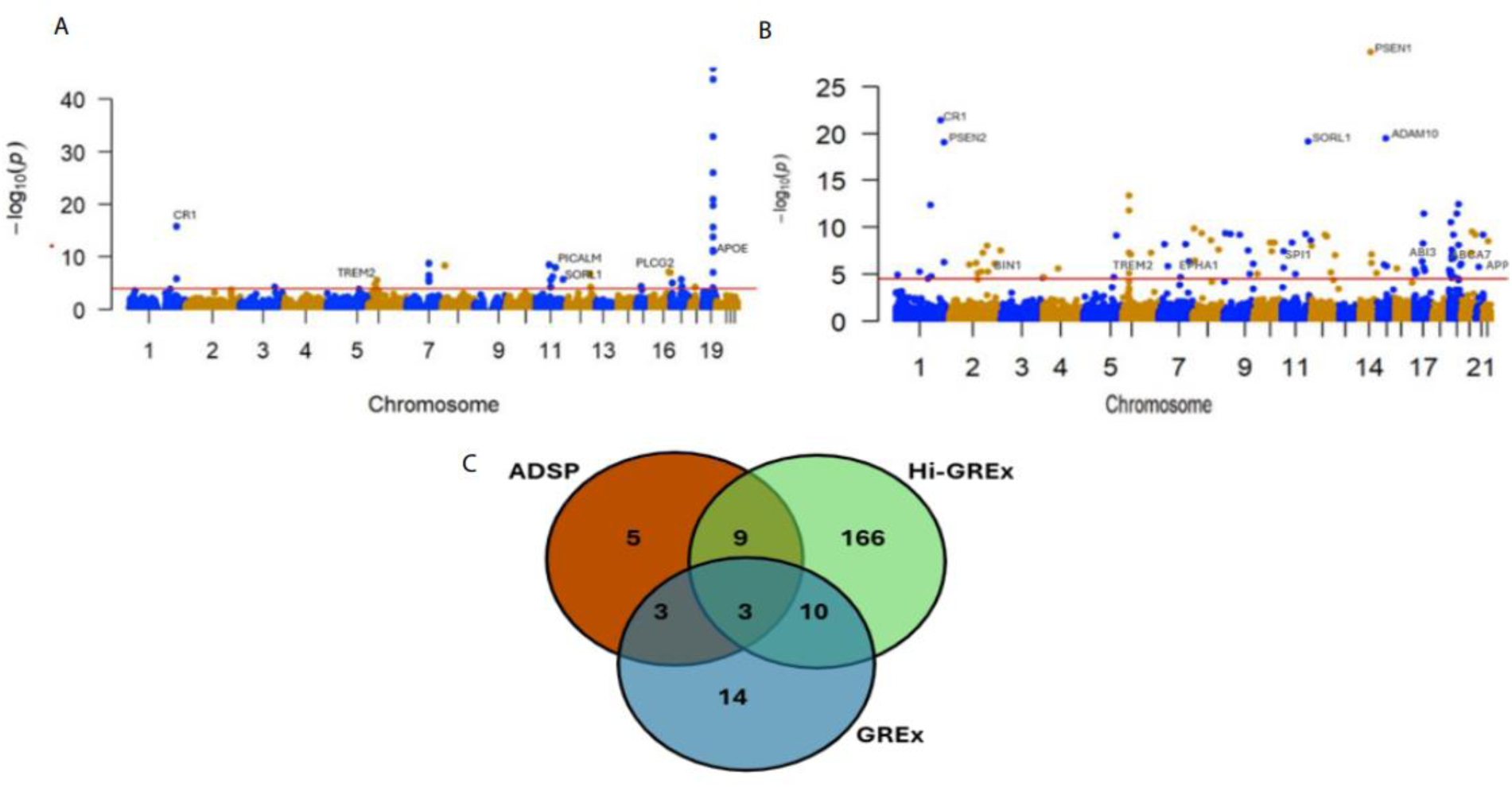
Manhattan plot of association results from the Alzheimer’s disease transcriptome-wide association study by utilizing (a) GREx models and (b): Hi-GREx models. The x-axis represents the genomic position of the corresponding gene, and the y-axis represents -log_10_-transformed association combined P value. Each dot represents the association for one specific gene. The labelled genes are significant genes which overlap with ADSP GVC data. (c): A Venn diagram demonstrates the overlap of significant gene hits across three datasets: ADSP-curated genes, GREx-based predictions, and Hi-GREx-based predictions

## Discussion

A major challenge in gene expression prediction modeling is to identify an optimal set of genetic features that yields maximal predictive accuracy for gene expression, particularly in cases where regulatory variants operate at varying genomic distances. Traditional approaches such as cis-based models (GREx) and models integrating long-range chromatin interaction information (Hi-GREx) each capture distinct facets of gene regulation,but may face limitations for specific loci. For instance, in the case of *APOE*, the Hi-GREx model’s reliance on predominantly long-range SNPs does not fully account for the gene’s regulatory complexity, as optimal predictive performance may require integration of both proximal and distal signals.

To address this, we introduced an alternative modeling strategy termed *sequential-GREx* (S-GREx) and Max_model [Supplementary_document.txt] [Figure S2]. Sequential integration acknowledges that certain genes may require a composite set of regulatory inputs to achieve optimal predictive fidelity. The maximal model (*Max-GREx*) for a given gene was defined as the model configuration yielding the highest spearman correlation coefficient among all candidates, thus operationalizing an empirical optimality criterion. This methodology acknowledges that the route to optimal gene expression prediction may involve multiple algorithmic paths, and it is only through systematic comparison and performance-based selection that the most effective strategy can be identified for downstream association analyses. Beyond the architecture of individual models, we recognize that the optimization landscape for gene expression prediction is inherently multifaceted. Similar to mathematical optimization, where a function may possess multiple local maxima and the global optimum is unknowable without broader exploration, gene expression modeling must empirically compare alternative feature selection strategies to identify the “best” model. Accordingly, for each gene, we developed multiple candidate models—cis-only (GREx), Hi-C guided (Hi-GREx), and sequentially integrated (S-GREx). The predictive performance of each model was evaluated by computing Spearman’s correlation coefficient (Rho) between the genetically predicted and residual measured expression values. 10563/13268 (79.6%) of the modelled genes show better performance in S-GREx compared (average Rho: 0.48) to GREx models [Figure S3A]. The Venn diagram llustrates the overlap and uniqueness of genes modeled by the three strategies (GREx, Hi-GREx, and S-GREx), revealing that while a large core set of 13,268 genes is captured by all methods, each model also identifies unique genes, reflecting the complementary nature of the regulatory features each approach incorporates [Figure S3B]. We compare performance of GREx, Hi-GREx, and S-GREx in terms of significant gene modeling across increasing Spearman correlation (ρ) thresholds. At relaxed thresholds, GREx captures the largest number of genes overall and contributes more unique model than either Hi-GREx or S-GREx. However, as the correlation threshold becomes more stringent, Hi-GREx retains a substantially larger fraction of significant models, underscoring its robustness and capacity to identify high-confidence predictions. This pattern highlights that while GREx is more inclusive at lower thresholds, Hi-GREx achieves superior predictive accuracy under stricter criteria. In pairwise comparisons, the combination of Hi-GREx and S-GREx uniquely recovers a greater number of genes compared to GREx-based pairs, suggesting that S-GREx contributes complementary regulatory information that synergizes most effectively with Hi-GREx. Together, these results indicate that Hi-GREx not only delivers the most reliable high-confidence models but also benefits from integration with S-GREx to expand gene coverage, thereby offering both accuracy and broader discovery potential. [Figure S3C]. The range of spearman correlation is higher for Hi-GREx modelled gene and we can achieve a maximum correlation of 0.91 when incorporating Hi-C selected long-distance SNPs in comparison to just S-GREx or GREx models [Table S1]. Together, these results emphasize the heterogeneous nature of gene regulation and the value of combining multiple modeling strategies to maximize gene expression prediction accuracy.

S-GREx still shows limited performance in identifying Alzheimer’s disease (AD)-associated genes. Specifically, S-GREx identified 96 significant genes and showed relatively low overlap with the established ADSP risk gene list in comparison to Hi-GREx models [Figure S4A, Figure S5]. This suggests that although S-GREx attempts to integrate long-range chromatin interactions, its predictive power and sensitivity remain suboptimal. In contrast, the Max-GREx model, which selects the best-performing model per gene across all strategies identified 216 significant AD-associated genes [S-PrediXcan_AssociationResults.xlsx] [Figure S4B]. It also showed greater overlap with the ADSP gene list, indicating that aggregating information across models, including Hi-C–informed ones, enhances both the detection of novel associations and the recovery of known AD risk loci [Figure S5]. Overall, these findings highlight that while S-GREx offers an alternative modeling approach, the Hi-GREx model provides superior power and biological relevance for AD gene discovery.

To elucidate the biological relevance of these findings, we performed gene set enrichment analysis on the significant brain tissue–associated genes. Consistent with previous literature implicating endocytic pathways in amyloid protein degradation and AD progression, we observed enrichment of 13 genes in the endocytosis Gene Ontology category—of which LRRTM1, SMAP1, and RUBCN, though absent from the ADSP gene list, have been independently implicated in AD risk in other studies [29,30]. Additionally, nine AD risk genes identified in this analysis (including PSEN2, SORL1, BIN1, ABCA7, ADAM10, PSEN1, CLU, PICALM, and APOE) were enriched in the amyloid precursor protein catabolic process (P = 4.94 × 10⁻ ⁸), underscoring the mechanistic relevance of these integrative models. Immune-related pathways were also highly significant, with enrichment in glial cell activation (P = 3.50 × 10⁻ ⁶), leukocyte-mediated immunity (P = 2.77 × 10⁻ ¹¹), and MHC protein complex assembly (P = 2.29 × 10⁻ ⁵) (Supplementary Table S3-Gene_Enrichment.txt). These results highlight a prominent role for immune function–related genetic pathways in the etiology of AD.

Our study demonstrates that incorporating Hi-C–derived long-distance SNPs dramatically improves gene expression prediction and the discovery of Alzheimer’s disease–associated genes, outperforming traditional gene-based models. The Hi-GREx approach not only expands the pool of informative SNPs, enabling detection of additional risk genes, but also enhances biological interpretability by capturing key pathways in AD pathogenesis. While alternative modeling strategies allow for flexibility in achieving optimal predictions, integrating distal regulatory information from Hi-C data is fundamental for maximizing sensitivity and insight in transcriptome-wide association studies. Future efforts should focus on extending this framework to multi-tissue studies, which will require Hi-C data across diverse tissues to uncover shared and tissue-specific regulatory mechanisms, providing a more comprehensive understanding of gene regulation and disease etiology.

## Method

### 1. Data Processing and Quality Control

#### Genotype and Expression Data Preparation

To ensure methodological consistency, we processed RNA sequencing and genotype data from GTEx brain cortex samples following the pipeline established by PrediXcan [7]. Whole genome sequencing data for brain cortex (n = 205) were downloaded from dbGaP in VCF format. Genotype missingness was addressed via the Michigan Imputation Server [20], leveraging the EAGLE phasing tool and the 1000 Genomes Phase 3 (GRCh38/hg38) reference panel, limited to European ancestry samples. Post-QC, the input comprised ∼960K SNPs filtered for Hardy-Weinberg equilibrium (P > 0.05) and minor allele frequency (MAF > 0.05). Only imputed SNPs with R² > 0.8 and MAF ≥ 1% were retained for further analysis. dbSNP version 152 was used to map rsIDs to SNPs based on allele and genomic position. SNP genotypes were encoded on a 0–2 scale to reflect the estimated dosage of the effect allele.

#### Expression Data Processing

Gene expression data for 18,916 genes in the brain cortex were obtained from the GTEx portal [21]. Read counts were normalized to RPKM, and then further subjected to inverse quantile normalization. Genes with expression above 0.1 RPKM in at least 20% of samples were retained. To account for confounders and batch effects, we applied Peertool software [22] to estimate 30 PEER factors which is determined based on sample size, used as covariates in multiple linear regression. Regression residuals were extracted and used as expression traits for model training.

### 2. Hi-C Data Integration and Genomic Alignment

#### Brain Cortex Hi-C Data Acquisition and Processing

We retrieved 3D Hi-C chromatin interaction data from developing human cerebral cortex [23], mapped to the hg19 reference genome at 40Kb bin resolution. Using UCSC’s LiftOver tool [24], we converted Hi-C interactions to hg38 coordinates to ensure cross-data compatibility. Only high-quality, uniquely mapping paired-end reads were selected for downstream chromatin interaction analysis.

#### Hi-C selection to capture spatial long-distance SNPs

To avoid scanning the entire genome for distal SNPs, we are utilizing brain cortex Hi-C to capture the “informative” distal SNPs that have chromatin interactions with its distal gene region. We only considered chromatin interactions between SNP-gene pairing that belong to different bins in our analysis. Every SNP-gene pair examined in the GTEx research was mapped to a bin pair of 40 Kb in Hi-C data. We concentrated on bin pairs which include at least one tested gene; that is, a gene position to be tested should be present in a bin must have a complementary long distance SNP position in the other bin. The bin pairs in our method also had distance > 1Mb so that the genomic distance (D=|i −j|) between bin i and bin j should be more than 1Mb to account for distal SNP-gene pairs. The resulting SNP-gene pair that mapped to a bin pair with corresponding chromatin interaction greater than 0 were selected as “informative” long distance SNPs to be used as a predictor

### 3. Feature selection

#### Feature selection the informative SNPs based on eQTL

To ensure parsimony and avoid overfitting when modeling gene expression, we implemented a rigorous feature selection step based on MatrixeQTL association statistics [25]. For each gene, we considered a comprehensive pool of candidate SNPs, combining both proximal (cis; within 1 Mb of the gene) and Hi-C-selected long-distance (distal) SNPs. we rank the SNPs based on eQTL association pvalue to determine SNPs linked to alterations in the expression of gene and select top features as our final input SNPs.

### 4. Gene Expression Prediction Models

We modeled genetically regulated gene expression (GREx) in the human brain cortex using an additive predictive framework similar to PrediXcan [7]. For each gene, the genetic contribution to expression was estimated as a weighted sum of selected SNP genotypes.

#### Additive Modeling Framework

For gene (g), the predictive model takes the following general form:

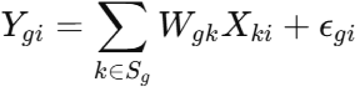

Where,

*Y_gi_* is the normalized expression of gene (g) in individual (i),

*X_ki_* is the genotype dosage (number of effect/reference alleles: 0, 1, or 2) for SNP (k) in individual (i),

*W_gk_* is the estimated effect size (weight) of SNP (k) on gene (g),

*S_g_* is the selected set of SNPs predicting gene (g),

*ε_gi_* is the residual term, capturing non-genetic and unmodeled effects.

The model is trained using elastic net regression, enabling robust variable selection and regularization in the high-dimensional SNP feature space.

#### Conventional GREx Model

In the standard GREx framework [Figure 1D-Panel A], only proximal (cis/short-distance) SNPs—those within ±1 Mb of the gene body (defined by GENCODE v26, GRCh38) [26]—are used as predictors:

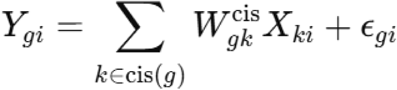

Where *cis(g)* denotes the set of all SNPs within 1 Mb of gene (g). This captures local, short-range regulatory contributions to gene expression

#### Hi-GREx Model

To capture both local and distal regulation, Hi-GREx incorporates not only proximal SNPs, but also distal SNPs selected based on chromatin interaction evidence from brain cortex Hi-C data to develop gene expression prediction model [Figure 1D-Panel B]:

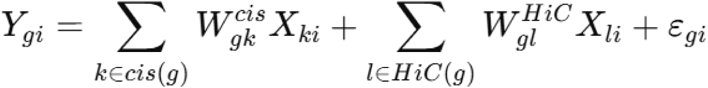

Where,

*cis(g*) denotes the set of all SNPs within 1 Mb of gene (g).

*HiC(g)* is the set of Hi-C selected, long-range distal SNPs shown to have physical contact with gene (g) in 3D space.

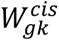 *and*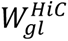 are the elastic net-estimated effect sizes for the cis and Hi-C distal SNPs, respectively

### 4. Transcriptome-Wide Association Analysis Using S-PrediXcan

Genetically regulated gene expression models like GREx and enhanced models incorporating 3D chromatin interaction data (Hi-GREx) were used for developing association with AD phenotype. S-PrediXcan [19] [Supplementary_document.txt] was then used to integrate GWAS data with gene expression prediction models and covariance matrices, yielding gene-level association statistics. Multiple testing correction was performed using the Benjamini-Hochberg false discovery rate (FDR), and genes surpassing the FDR threshold were considered significantly associated with AD risk. Associations were compared across cis-only and 3D-informed models to evaluate the added value of incorporating distal regulatory information.

## Supporting information

Results of Gene Ontology (GO) enrichment analysis showing significantly enriched biological processes and molecular functions for the significant gene

S-PrediXcan association results for significant genes across GReX, Hi-GReX, S-GReX, and comparative models.

Supplementary material presenting methodology details, performance evaluations, and association analyses of alternate strategies

## Availability of data and code

All the codes used to develop GREx, Hi-GREx and S-GREx gene expression prediction model are kept in the <u>HiGREx_Tutorial</u> Github repository with tutorial dataset for the user to try and develop models.

Other software and data URLs:

PrediXcan: <u>PrediXcan_Predictdb</u>, University of Michigan Imputation-Server: <u>Michigan_Imputation</u>, GTEx Portal: <u>GTEx,</u> S-PrediXcan software: <u>S-PrediXcan</u>, Brain-Cortex Hi-C (GEO: <u>GSE77565</u>), Peertool: <u>Peertool_Software</u>, MatrixeQTL tool: <u>MatrixeQTL</u>

## Author Information

## Ethics declaration

### Ethics approval and consent to participate

Not applicable.

### Consent for publication

Not applicable.

## Supplementary information

Supplementary_document.docx

Supplementary material presenting methodology details, performance evaluations, and association analyses of advanced gene expression prediction frameworks (S-GREx, Max-Model) in the context of Alzheimer’s disease transcriptome-wide association studies.

S-PrediXcan_AssociationResults.xlsx

Comparison_of_Performance_ADGenes.xlsx

Comparison of prediction performance across GReX, Hi-GReX, and S-GReX models for Alzheimer’s disease–related genes.

Supplementary_Table_S3-Gene_Enrichment.txt

Results of Gene Ontology (GO) enrichment analysis showing significantly enriched biological processes and molecular functions for the selected significant AD gene set.

