## Supplementary material presenting methodology details, performance evaluations, and association analyses of alternate strategies for "Hi-GREx: A 3D Genome–Guided Framework for enhancing Gene Expression Prediction Using Hi-C–Selected Distal SNPs"

**Supplementary Information**

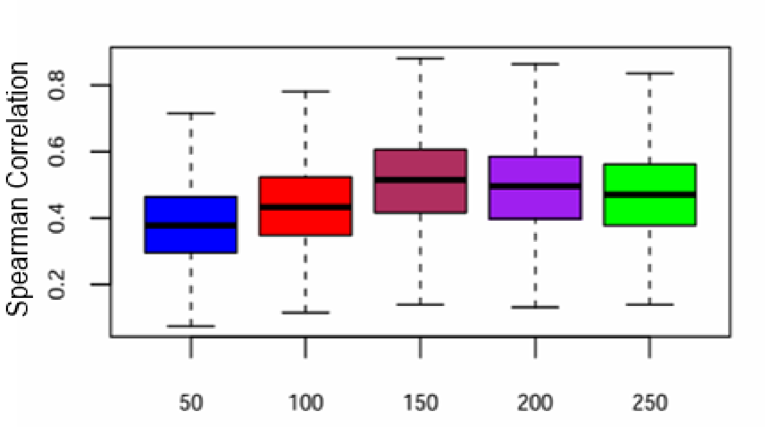


**Figure S1**: Boxplot showing the distribution of correlation across different “n” SNPs input features for genes across chromosomes:1,2,3,20,21,22 (number of modelled genes=3509). This figure overall depicts that 150 is the optimal number of SNP features to get a good prediction accuracy in comparison to other “n” features.

Model Performance for GREx vs Hi-GREx vs S-GREx

| **Group** | **Samples** | **Genes** | **Mean $R^2$** | **Range $R^2$** |
|---|---|---|---|---|
| Model Performance in GTEx | | | | |
| GREx | 205 | 14134 | 0.46 | 0.10-0.88 |
| Hi-GREx | 205 | 16825 | 0.53 | 0.11-0.91 |
| S-GREx | 205 | 17460 | 0.48 | 0.11-0.88 |

**TableS1: Model performance comparison for GREx, Hi-GREx, and S-GREx gene expression prediction frameworks in GTEx brain cortex tissue.**

Summary of the number of samples, total genes modeled, and predictive accuracy

(Mean $R^2$ and Range $R^2$) for each approach. Inclusion of Hi-C–guided long-range SNPs in Hi-GREx enhanced the average model fit (Mean $R^2$ = 0.53), compared to conventional local SNP-based GREx models (Mean $R^2$ = 0.46). The S-GREx sequential approach further broadened gene coverage, modeling 17,460 genes. These results highlight improved predictive power and gene coverage when integrating Hi-C–derived chromatin interaction data.

| SNP_ID | GWAS_ReportedGene | Model_Gene |
|---|---|---|
| rs204480 | CLPMT1 | ABCA7 |
| rs679515 | CR1 | PSEN2 |
| rs10498633 | SLC24A4 | PSEN1 |
| rs867611 | PICALM | SPI1 |

**TableS2:** Table showing the association between specific single nucleotide polymorphisms (SNPs) and their reported genes from Genome-Wide Association Studies (GWAS), alongside corresponding model genes used in analysis. Each row lists an SNP identifier, the nearest gene reported by GWAS, and the gene for which the

particular SNP was used in the model.

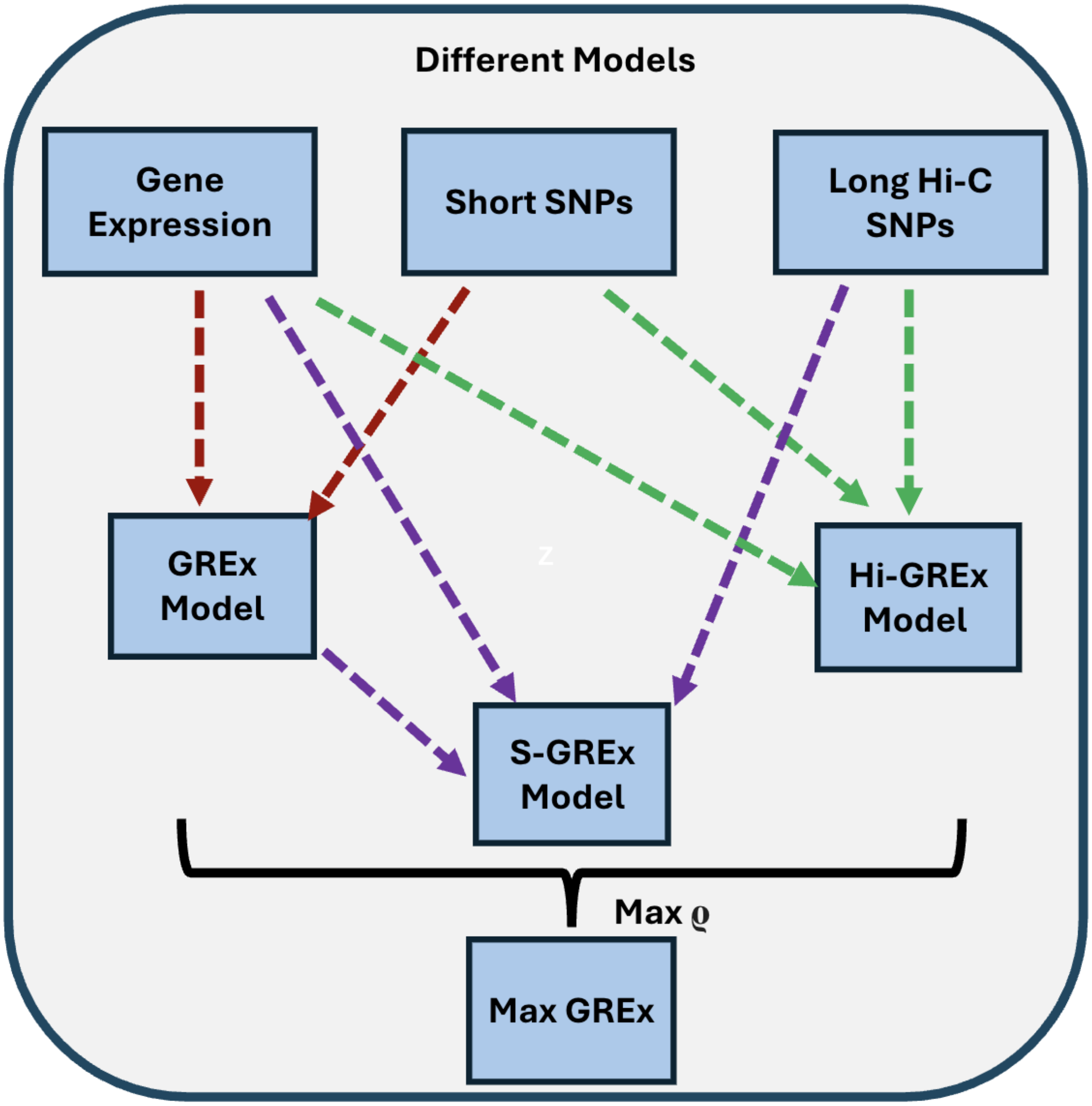


**FigureS2:** Schematic overview of different gene expression prediction models. Diagram showing how short SNPs (GREx), Hi-C long SNPs (Hi-GREx), and sequential models (S-GREx) each contribute to gene expression prediction. The Max-GREx approach selects the optimal model based on highest prediction performance.

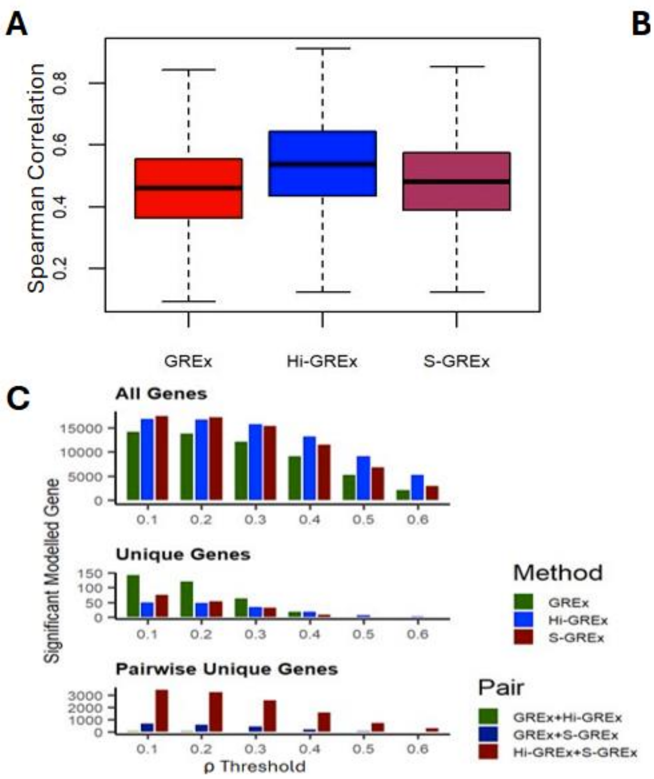


**FigureS3:** Comparative Evaluation of Gene Expression Prediction Models 3(a): Boxplots of Spearman correlation coefficients for three gene expression prediction models: *GREx*, *S-GREx*, and *Hi-GREx*. Each boxplot summarizes the distribution of correlations between predicted and observed gene expression across all genes for each model, illustrating overall predictive accuracy. As shown in Panel A, the *Hi-GREx* model yields higher median Spearman correlations compared to both *GREx* and *S-GREx*, indicating improved overall predictive accuracy.  (b): Venn diagram representing the overlap in genes successfully predicted by each model. The Venn diagram reveals the extent of overlap and uniqueness in gene coverage among the models: while a substantial set of genes is shared between models, each approach also predicts a

unique subset of genes not captured by the others. (c) Bar plots show the number of significant gene models identified by GREx, Hi-GREx, and S-GREx across different Spearman correlation (ρ) thresholds. The top panel (“All Genes”) shows total significant models per method; the middle panel (“Unique Genes”) highlights genes exclusively predicted by a single method; and the bottom panel (“Pairwise Unique Genes”) shows genes jointly predicted by two methods but absent in the third. Hi-GREx consistently identifies more significant genes across thresholds, reflecting its improved predictive performance.

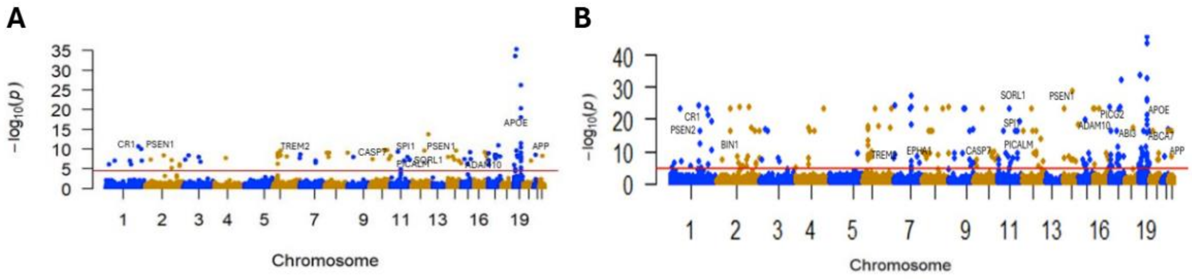


**FigureS4:** Manhattan plot of association results from the Alzheimer’s disease transcriptome-wide association study by utilizing (a) S-GREx models and (b): Max_models. The *x*-axis represents the genomic position of the corresponding gene, and the *y*-axis represents -log$_{10}$-transformed association combined *P* value. Each dot represents the association for one specific gene. The labelled genes are significant genes which overlap with ADSP GVC data.

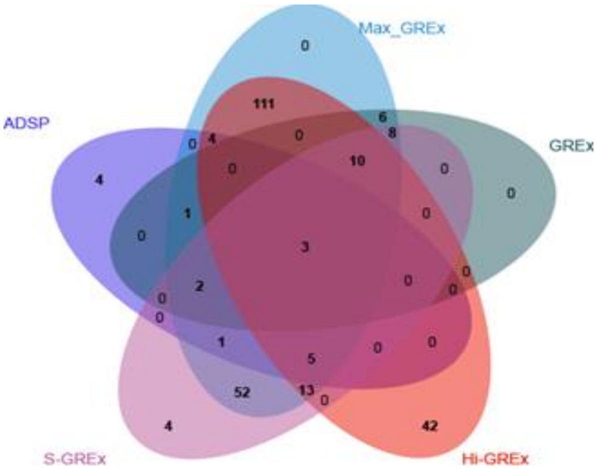


**Figure S5:** Venn diagram showing overlap between different significant genes across different models with well-known AD genes from ADSP GVC data

**Methodology for S-GREx, Max_Model and S-PrediXcan association study:**

1. **Sequential (S-GREx) and Maximal Gene Regulatory Expression Model Selection:**

   (i) S-GREx method involves two steps to generate the final prediction model. We first constructed a gene expression prediction model using only short-range (local) SNPs within the cis regulatory region. Subsequently, we augmented this model by sequentially incorporating long-range SNPs prioritized by chromatin interaction (Hi-C) data and ranked by eQTL signal, thereby resulting in an enriched predictor set encompassing both local and distal regulatory variants.

The S-GREx model (4) for gene (g  in individual (i) is defined as:

$$Y_{gi} = \sum_{k \in \text{cis}(g)} W_{gk}^{\text{cis}} X_{ki} + \sum_{m \in \text{seq}(g)} W_{gm}^{\text{seq}} X_{mi} + \epsilon_{gi}$$

*Where,*

- *cis(g)* is the set of SNPs within ±1 Mb of gene (g) (local regulatory region), used in the first model step [GREx]
- *seq(g)* is the set of Hi-C prioritized, distal SNPs selected and ranked by eQTL signal, sequentially added to the model in the second step
- $W_{gk}^{\text{cis}}$, $W_{gm}^{\text{seq}}$ are the elastic net-estimated effect sizes for the cis and sequenced (distal) SNPs respectively, $X_{ki}$ and $X_{mi}$ are genotype dosages for individual (i),
- $\varepsilon_{gi}$ is the error term for gene (g) in individual (i).

(ii) Max-GREx (maximal model): Once the model is developed in each case, we then evaluate the performance of the model by calculating Spearman's correlation coefficient between predicted expression and residual expression values across the individuals. For each gene, we select a maximal model which has the highest correlation among others for further association.

$$\text{Maximal\_Model} = \max\left(\rho\text{Grex},\ \rho\text{Hi-Grex},\ \rho\text{S-Grex}\right)$$

### 2. Transcriptome-Wide Association Analysis Using S-PrediXcan

We performed transcriptome-wide association studies (TWAS) to identify genes whose genetically predicted expression levels are associated with Alzheimer's disease (AD), leveraging the S-PrediXcan framework.

#### (i) Preparation of Predicted Expression Models and Reference Data

Genetically regulated gene expression prediction models were obtained from *PredictDB*, including both traditional models based solely on local (cis) regulatory variation, and enhanced models incorporating three-dimensional chromatin interaction data (Hi-C). These models estimate gene expression using genome-wide SNP data, accounting for both proximal and distal effects where appropriate. For each gene, covariance matrices of the SNPs included in the prediction model were prepared, as required by S-PrediXcan, to accurately model LD structure during association testing.

#### (ii) Genome-Wide Association Summary Statistics (GWAS)

GWAS summary statistics for Alzheimer's disease were curated and formatted according to S-PrediXcan requirements. Input files were provided in plain text or gzipcompressed formats, with customizable column mappings specified via command-line arguments.

**(iii) TWAS Procedure and Statistical Analysis:**

S-PrediXcan was executed for each gene and tissue model, integrating GWAS summary data, gene expression prediction weights, and SNP covariance matrices. The pipeline computes a gene-level association statistic, representing the correlation between genetically predicted gene expression and AD risk. Statistically significant associations were determined following multiple testing correction using the false discovery rate (FDR, Benjamini-Hochberg procedure).

Genes surpassing the FDR threshold were considered putatively implicated in AD risk. The resulting associations were further examined to compare models that incorporate three-dimensional genomic regulatory information against those capturing only local variant effects.
